# Pre-treatments of Ex vivo Vascularized Composite Allografts: A Scoping Review

**DOI:** 10.1101/2024.11.20.624373

**Authors:** Caroline E Baker, Thor S Stead, Alay R Shah, Dominika Pullmann, Sachin Chinta, David L Tran, Hilliard T Brydges, Matteo Laspro, Bruce E Gelb, Eduardo D Rodriguez, Piul S Rabbani

## Abstract

**PURPOSE:** The various physiological profiles comprising vascularized composite allografts (VCAs) pose unique challenges to preservation. Minimizing ischemia, reperfusion injury, and rejection remains a primary focus of graft pre-treatments (PTs). Currently, the gold standard PT consists of flushing the graft and placing it in static cold storage in the University of Wisconsin solution. With this method, graft viability is limited to four to six hours. Prolonging this time limit will increase donor allocation radius, access to care, and positive patient outcomes. We aimed to evaluate novel PTs that could potentially enhance and lengthen VCA viability.

**METHODS:** Following PRISMA guidelines, we conducted a comprehensive literature search of Embase, Cochrane, and PubMed. Studies had to be published prior to June 15, 2022. PTs had to target cell physiology, rather than immunogenicity. We extracted data including study design, PT details, evaluation metrics, and outcomes.

**RESULTS:** We identified thirteen studies, categorized into three groups: solution-based alterations to the gold standard, ex vivo perfusion, and other novel techniques. The incorporation of hydrogen sulfide and Perfadex as solutions in the gold standard protocol demonstrated a six-day delay in rejection and limited reperfusion injury markers, respectively. In one ex vivo perfusion study, after 24 hours of PT and 12 hours post-transplant, VCA muscle contractility remained close to normal. The gold standard PT did not demonstrate the same success. However, graft weight gain, up to 50% of baseline among the articles reviewed, is a prominent side effect of perfusion. Another technique, cryopreservation, displayed 90% graft failure by venous thrombosis, despite high free graft viability following two weeks of storage.

**CONCLUSIONS:** This study of pre-treatment modalities found a variety of encouraging preservation techniques for grafts with high levels of tissue diversity. Ex vivo perfusion dominated PT innovation with promising results in preserving the viability and functionality of muscle, which is central to the restoration of movement. Future studies are necessary to evaluate long-term graft outcomes and to optimize PT protocols for extended preservation times to ensure clinical relevance.

## Introduction

Vascularized composite allotransplantation (VCA) is the concurrent transplantation of multiple tissue types, including skin, blood vessels, muscles, and nerves, and offers an innovative surgical solution to congenital, traumatic, oncologic, and other defects not otherwise amendable to conventional reconstructive methods.^1,2^ Upper extremity, uterine, and facial grafts among others have been shown to confer significant functional recovery in patients with lifestyle-limiting defects.^3^ In 2014, the Organ Procurement and Transplantation Network officially classified vascularized composite allografts (VCAs) as “organs” that may be sourced from brain-dead human donors.^1^ Over the next seven years, 22 medical centers performed 64 VCA transplantations.^2,3^ However, the heterogeneous tissue architecture of these grafts creates unique barriers to their widespread integration into transplant medicine. The constituent tissue types differ considerably as they each feature distinct homeostatic and recovery mechanisms.^4–6^ Additional considerations of VCA include immune compatibility between donors and recipients, the antigenicity of skin, the continuous exposure of the transplanted graft to the external environment, and the potential functional recovery of muscle and nerves.

A key opportunity for promoting safe and successful VCA transplantation is the time interval between procurement and transplantation. During this time, ischemia becomes a key variable in determining the extent of injury to the graft and potentially affecting graft survival.^6^ Once removed from the donor, energy stores are quickly depleted, inhibiting mitochondrial activity, and damaging DNA. Subsequent signaling cascades lead to cellular apoptosis and the release of harmful metabolic by-products including reactive oxygen species (ROS) and proinflammatory cytokines.^7^ Additionally, re-anastomosis upon transplantation introduces the potential for ischemia reperfusion injury (IRI), which may contribute to the risk of graft failure.^6,8–10^ The existing VCA literature purports that graft optimization during this *ex vivo* ischemic period will decrease the risk of rejection and promote successful VCA transplantation. Studies in solid organ transplantation (SOT) demonstrate a critical role of pre-treatments (PTs), which may be applicable in prolonging VCA viability as well.^11^

The current gold standard PT for SOT consists of flushing the graft with the University of Wisconsin (UW) solution to remove remaining blood products and metabolic waste.^12^ The graft is then kept in static cold storage (SCS), usually at 4°C in UW solution, to slow cellular metabolism and delay ischemic injury.^7,11,12^ UW solution is a low-sodium, high-potassium solution that demonstrated early success in SOT and among many functions, protects mitochondrial membrane integrity, preserves energy stores, and counteracts hypothermia-induced cellular swelling.^11,13–15^ Evidence of viable tissue pre-transplantation and preserved function post-transplantation justified the widespread adaptation of UW solution to safely extend cold ischemia time up to 36 hours for kidneys and 6 hours for hearts and lungs.^13,14,17^ Low early complication rates and increased long-term graft survival further support its widespread use in *ex vivo* solid organ static preservation.^13,14^

In contrast, current standards for VCA preservation in UW solution limit cold ischemia time to 4 to 6 hours. This interval restricts the allowable geographical distance between donors and recipients and contributes to a relatively small donor pool highlighting the need to maximize the preservation interval.^16^ In this scoping review of the literature, we aim to study existing VCA PT strategies and their effects on graft viability. Our analysis indicates the need for the development of PTs specifically for VCA to enhance graft survival and optimize functional recovery.

## Methods

Following Preferred Reporting Items for Systematic Reviews and Meta-Analyses (PRISMA) guidelines, we conducted a literature search of PubMed, Cochrane, and Embase using 33 terms related to VCA and transplantation (**Supplemental Table 1)**.

Two authors (TSS and ML) conducted the literature screening. A third author (HTB) resolved conflicts between reviewers. Articles which studied PT methods on *ex vivo* VCA and which were published prior to June 15, 2022 were included. Additionally, PT interventions had to target cell physiology, rather than immunogenicity. Criteria for exclusion included abstracts, commentaries, editorials, non-English articles, and review papers or other articles not reporting primary data.

Following screening, another author (CEB) extracted data regarding study design, PT characterization (process, solutions, timing, etc.), evaluation metrics, and outcomes. Studies were then categorized one of three ways based on the pre-treatment method: solution-based alterations to the gold standard (static cold storage [SCS] in UW solution), *ex vivo* perfusion, or other techniques.

## Results

We identified 4,999 studies including 3,125 unique articles. After screening, 13 studies describing 12 distinct methods of PT were included for data extraction. The exclusion breakdown is depicted in **Figure 1**. *Ex vivo* perfusion was the most commonly studied method, followed by other novel techniques with four, and solution-based pre-treatments with two. We summarize the 13 articles within their categorical distinctions and their characteristics in **Supplemental Table 2**.

**Figure 1.**
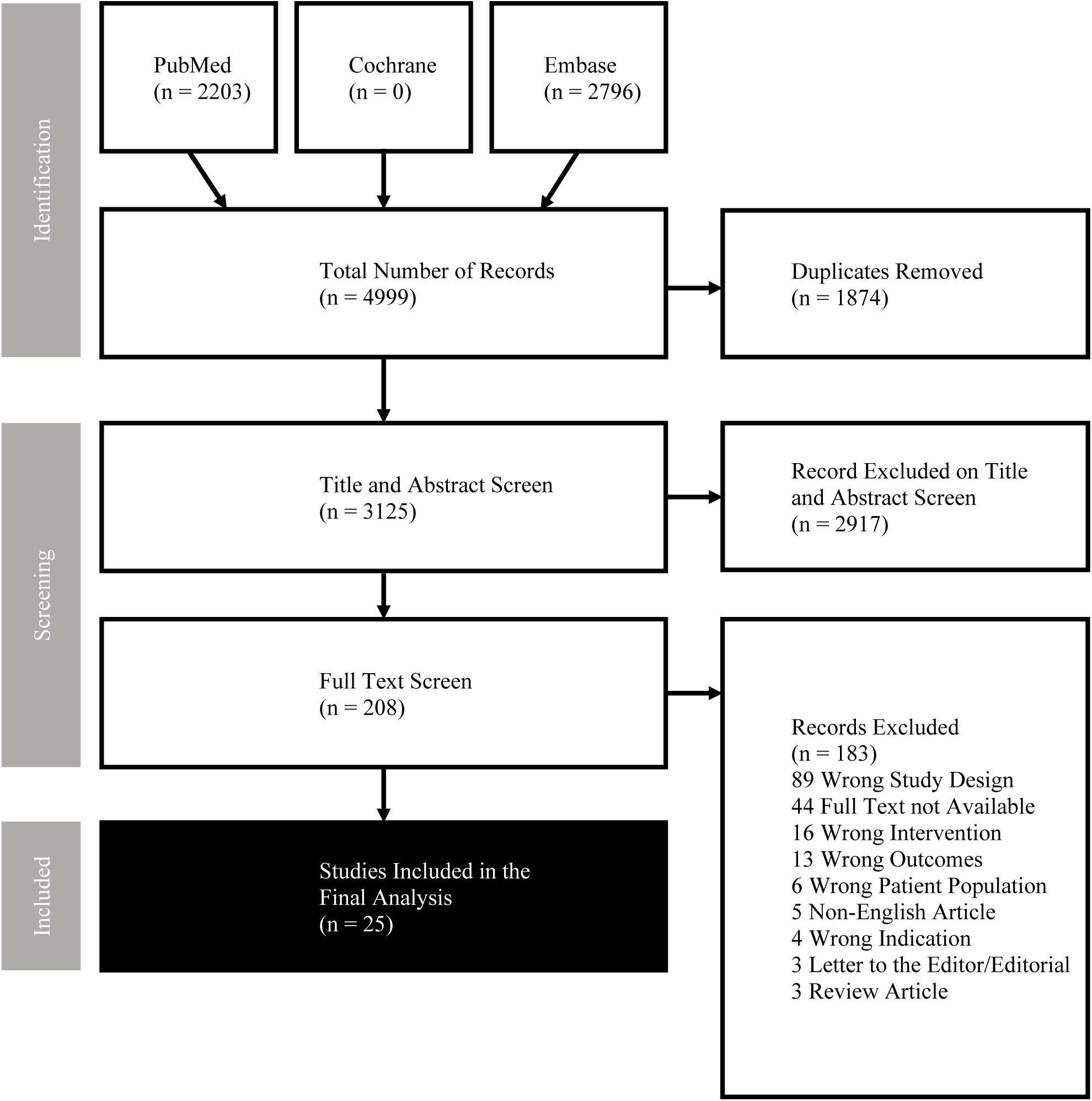
Exclusion breakdown for article screening.

The studies were primarily preclinical mostly utilizing porcine and murine models and a canine model, but also included one study on human subjects. Researchers performed allogeneic transplants, syngeneic transplants, or replants or opted solely to analyze graft viability without transplantation. Studies used limb grafts, musculocutaneous flaps, and superficial flaps. All grafts contained skin, blood vessels, muscle, subcutaneous/adipose tissue, and fascia, and a minority of studies also surgically repaired nerve, muscle, tendon, and bone to analyze recovery of graft movement.^12,18–29^ **Figure 2** highlights notable advantages and disadvantages of each approach.

**Figure 2.**
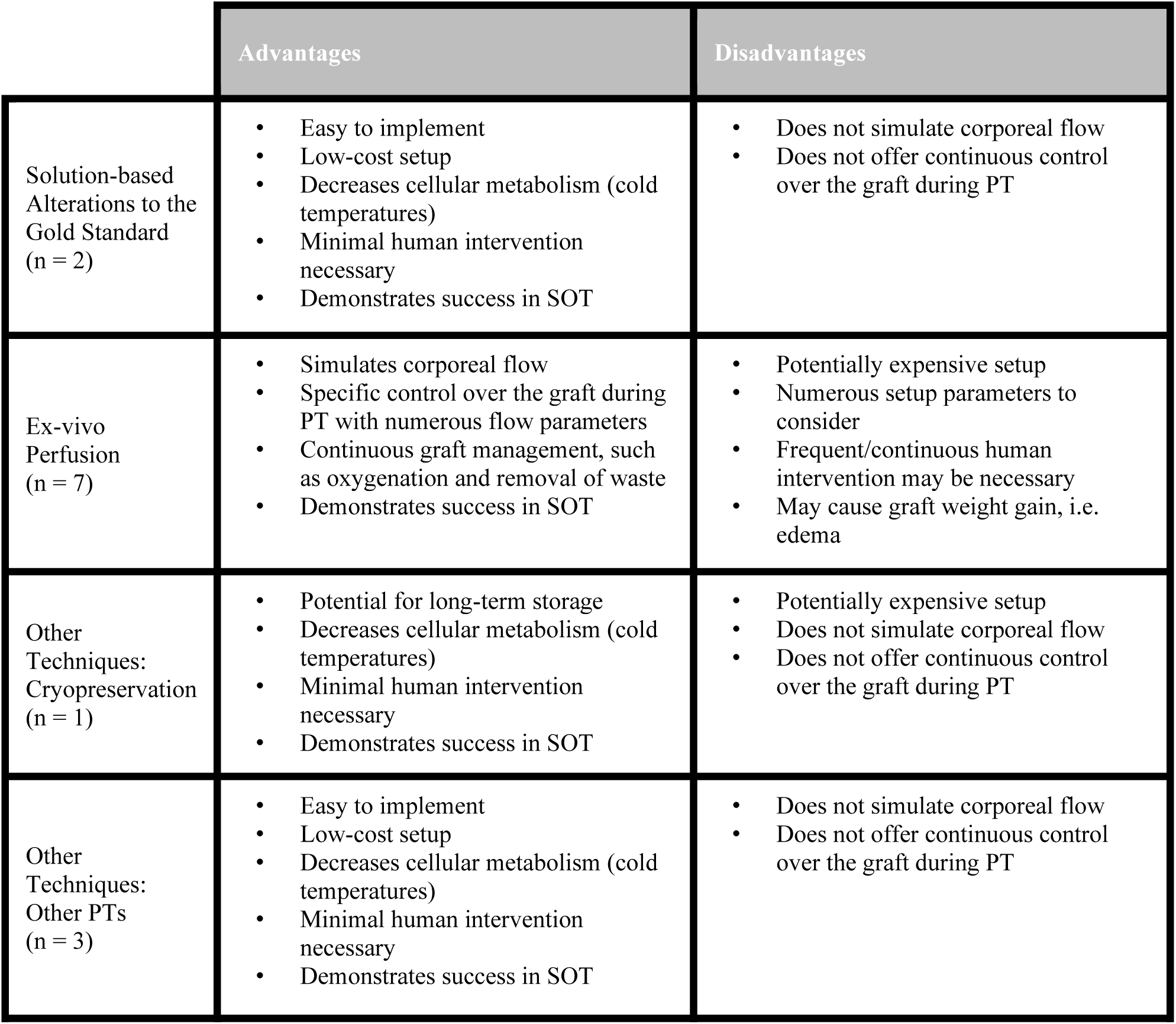
Advantages and disadvantages of PT methodology categories. SOT = solid organ transplantation; PT = pre-treatment.

### Solution-based Alterations to the Gold Standard

Two studies investigated the addition of two different solutions to UW solution both at the time of flushing a newly harvested graft and during SCS. Additive solutions studied included hydrogen sulfide (H_2_S) and histidine-tryptophan-ketoglutarate (HTK), while Perfadex was studied as an additional currently accepted preservation solution. One study assessed the effect on acute rejection examining graft erythema, edema, and gross necrosis as markers of clinical rejection. Histological analysis consisted of hematoxylin and eosin (H&E) staining and CD3 staining to indicate T-lymphocyte infiltration. Another study evaluated ischemia-reperfusion injury (IRI) by measuring apoptotic markers such as cleaved caspase 3 (CC3) and by employing the terminal deoxynucleotidyl transferase dUTP nick-end labeling (TUNEL) assay. IRI was also assessed by inflammation-associated markers, such as high-mobility group box 1 (HMGB-1) and laminin, a marker of tissue integrity. **Table 1** provides a comparison between the study designs.

**Table 1.**
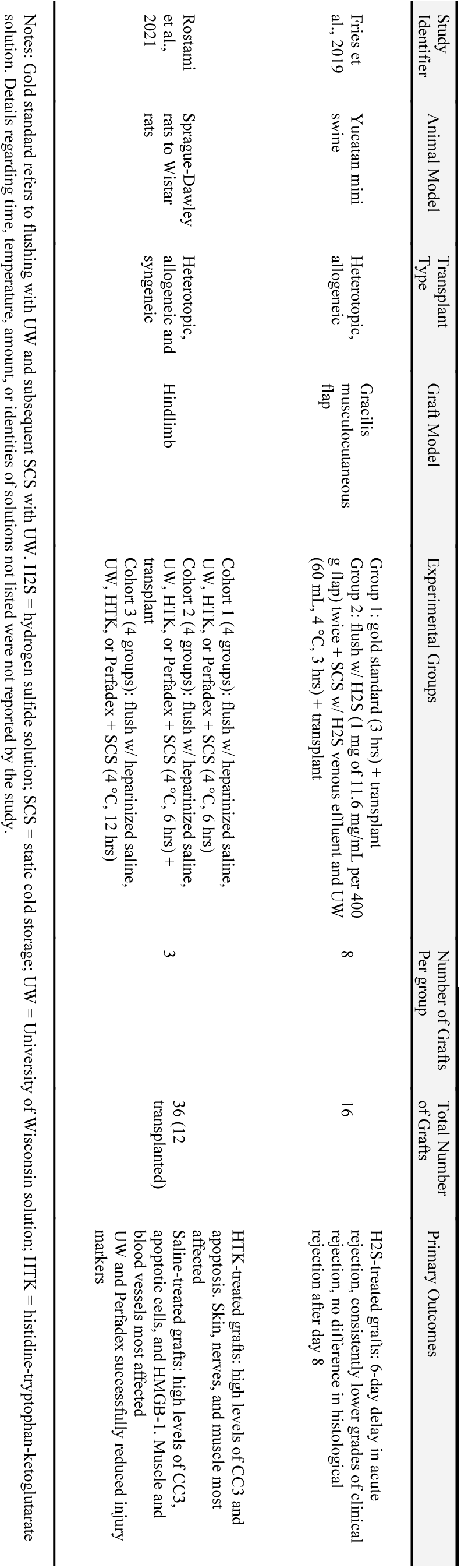
Characterizations of experimental solution-based alterations to the gold standard VCA pre-treatment protocol.

Fries et al reported a 6-day delay in the onset and degree of acute rejection with H_2_S-treated grafts when compared to UW solution alone. In contrast, compared to clinical rejection, histological grading of rejection was found to be more severe, though there was no statistically significant difference between groups after day 8.^18^

Rostami et al observed increased injury markers with increasing SCS time, both in allogeneic vs. syngeneic transplantation as well as before and after transplantation. There was an increase in apoptotic cells within the skin, muscle, and nerves noted at 12 hours of SCS in HTK compared to other solutions. The saline-treated group showed higher levels of CC3, apoptotic cells, and HMGB-1 in the muscle and vasculature when compared to UW solution and Perfadex groups. There were no detectable differences in injury markers for bone across any of the solutions. Overall, the success in limiting injury markers made UW and Perfadex preferred preservation solutions in this study.^19^

### *Ex vivo* Perfusion

Mimicking corporeal flow dynamics, *ex vivo* perfusion is the continuous passage of fluid through the donor graft vasculature, prior to transplantation. Flow parameters including pressure, rate, temperature, and duration of perfusion varied across studies. ^20,21,24,26–29^ Most of the perfusion studies reviewed incorporated a pre-perfusion flush.^20,21,26,27,29^ Reported perfusates included oxygenated blood-based or acellular solutions, including UW, HTK, skeletal muscle media, and Steen solution, which is another accepted preservation solution used in lung transplantation. Additives to the perfusates included heparin to prevent clotting, dextrose/dextran to maintain osmotic balance, steroids to minimize inflammation, insulin to control potassium levels, and glucose for cellular metabolism.^20,21,24,26–29^ Some studies employed periodic replacement of perfusate to mitigate toxic waste buildup.^20,21,27^ To evaluate graft viability, studies conducted clinical assessments of graft skin, histological assessments including TUNEL assays, and assessed myoglobin and creatine kinase levels as markers of muscle injury. Gene expression analysis included *hemoglobin-based oxygen carrier-201 (HBOC-201)*, *hypoxia-inducible factor 1-alpha (HIF1α)*, *tumor necrosis factor receptor superfamily member 10-A (TNF-SR-10A)*, *regulator of G-protein signaling 2 (RGS-2)*, *nuclear factor kappa beta inhibitor zeta (NFκBIZ)*, *interleukin-1 beta (ILβ1)*, *fibroblast growth factor 6 (FGF6*), *DNA damage-inducible transcript 4 (DDIT-4)*, and *caspase-3 (Casp3*).^20,21,24,26–29^ Studies also examined graft function by measuring muscle contractility to electrical stimulation and force generation and also measured metabolic profiles to evaluate organ and cell level function.^20,21,24,27,29^ **Table 2** compares the seven study designs.

**Table 2.**
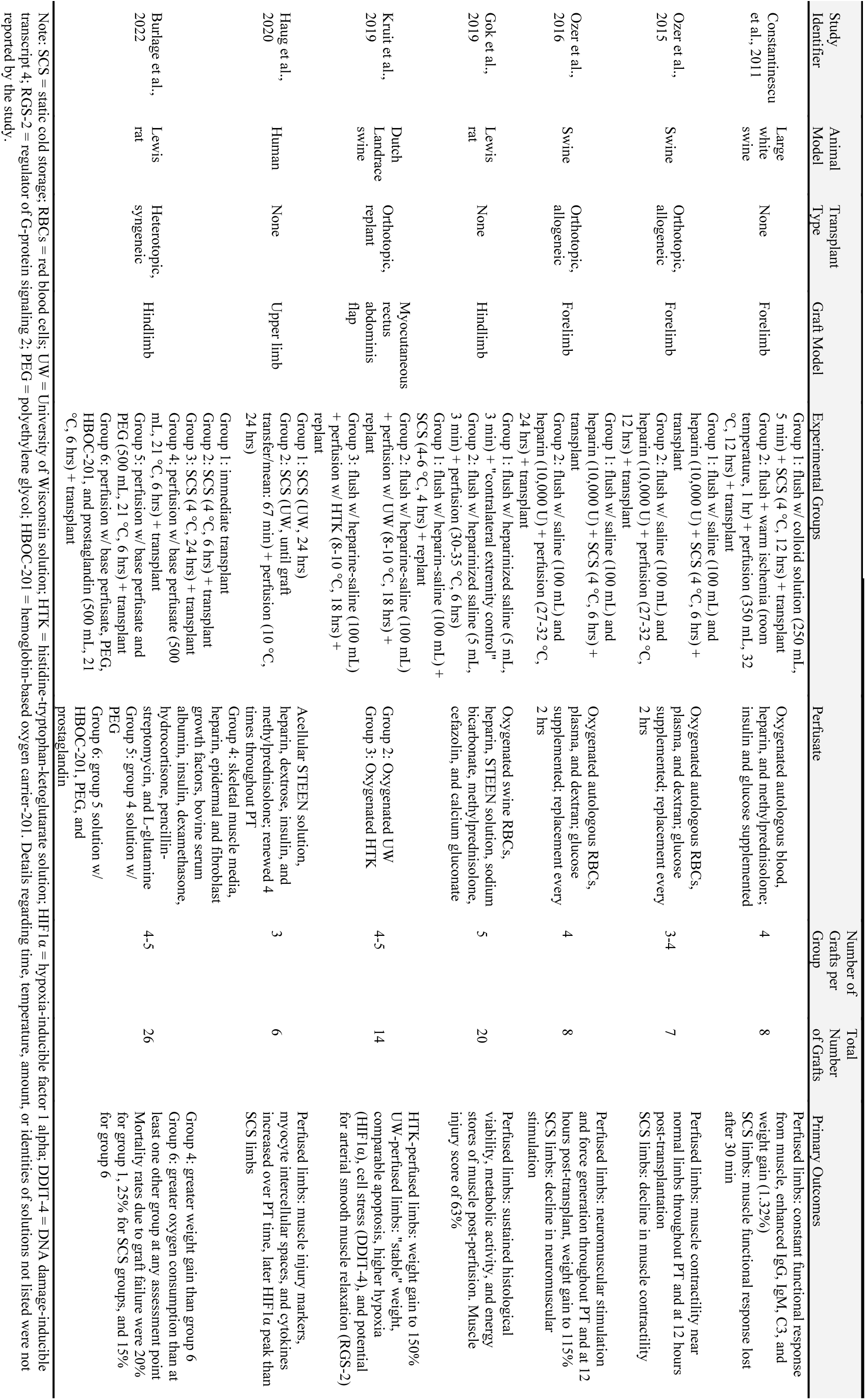
Characterizations of experimental ex vivo perfusion pre-treatment protocols.

When comparing perfusion to SCS, Haug et al found slightly lower overall levels of HIF1α and a 12-hour delay in peak levels (post-operative hour 8 vs. 20).^28^ During the IRI interval, Kruit et al confirmed differential expression of apoptotic factors, markers of ischemia, pro- and anti-inflammatory markers, muscle injury markers, and markers of cellular stress response. Compared to HTK-perfused flaps, UW-perfused flaps exhibited greater hypoxia and cell stress, comparable apoptosis, but better endothelial protection.^26^ In a study by Burlage et al, both polyethylene glycol (PEG) and prostaglandin-supplemented skeletal muscle media and supplemented HBOC-201 solution, showed decreased rates of graft failure when compared with SCS or even immediate transplantation.^24^ All studies showed evidence of weight gain in perfused flaps over SCS controls while lactate and potassium levels were well-controlled.^24,26,28^

Studies examining muscle contractility and force generation found preservation of muscle function in up to 24 hours of *ex vivo* perfusion when compared to SCS.^20,21^ Ozer et al 2015 also demonstrated single-muscle fiber contractility comparable to non-operated limbs.^20^ Notably, this effect was diminished after 18 hours of perfusion in a follow-up Ozer et al 2016 study but also demonstrated recovery within 24 hours following transplantation.^21^ In contrast, SCS showed a gradual decline in muscle contractility and force generation.^20,21,27^ Perfused grafts were found to have metabolic activity and energy stores comparable to baseline.^29^ However, perfused muscle showed a significantly greater injury score (62.87 ± 5.69% vs. 4.12 ± 3.12%), increased immunoglobulins and complement expression, and increased muscle injury markers compared to SCS controls.^27–29^

### Other Techniques

Four studies were included for review investigating modalities not otherwise specified in the literature such as cryopreservation of grafts at –140°C, post-SCS flush with lactated Ringer’s solution to remove leukocytes and decrease graft antigenicity, and a post-harvest flush. To evaluate graft viability, studies used clinical assessments of graft skin, histological evaluation including H&E stains and a factor VIII stain, metabolic profiles, and inflammatory profiles measured by flow cytometry.^12,22^ Compound muscle action potential (CMAP) amplitude, latency, and other electromyography evaluations were completed to evaluate graft muscle and nerve function as well.^23,25^ A summary of the four study protocols is shown in **Table 3**.

**Table 3.** Characterizations of other PT techniques.

| Study Identifier | Animal Model | Transplant Type | Graft Model | Experimental Groups | Number of Grafts per Group | Total Number of Grafts | Primary Outcomes |
| --- | --- | --- | --- | --- | --- | --- | --- |
| Rinker et al., 2008 | Lewis rat | Orthotopic, syngeneic | Epigastric flap | Group 1: immediate transplant<br>Group 2: perfusion w/ 2.0 DMSO + transplant<br>Group 3: flush w/ RL and UW + SCS w/ RL and UW (4 °C) + cryopreservation w/ 2.0 DMSO (-140 °C, 2 weeks) | 10 | 30 | Group 1: 100% of grafts survived to 60 days<br>Group 2: all but one graft survived to 60 days<br>Group 3: post-preservation viability of 91.9% and no difference in epithelial viability index from fresh grafts. Only one graft survived to 60 days. |
| Liu et al., 2010 | Beagle canine | Orthotopic, replant | Hemifacial flap | Group 1: gold Standard (4 °C, 24 hrs) + flush w/ RL (500 mL) + transplant<br>Group 2: gold Standard (4 °C, 36 hrs) + flush w/ RL (500 mL) + transplant<br>Group 3: gold Standard (4 °C, 48 hrs) + flush w/ RL (500 mL) + transplant | 2-3 | 8 | Muscle and nerve viability decreased most quickly, skin the least<br>Decreased muscle function from baseline<br>Statistically significantly higher graft survival for the 24-hour group over 36 and 48 hours. A preservation time limit of 24 hours is recommended. |
| Amin et al., 2018 | Swine | None | Forelimb | Group 1: flush w/ UW (500 mL) and heparin (10,000 U) + SCS (2 hrs) + flush w/ RL<br>Group 2: flush w/ UW (500 mL) and heparin (10,000 U) + SCS (6 hrs) + flush w/ RL | 6 | 12 | The leukocyte population was dominated by T cells and lacked anti-inflammatory cytokines<br>The only significant difference in leukocytes between the flush at 6 hours of SCS and 2 hours was a decrease in CXCL-8 |
| Gok et al., 2020 | Lewis rat | Orthotopic, syngeneic | Hindlimb | Group 1: naive control<br>Group 2: sciatic nerve transection + repair (proof-of-concept)<br>Group 3: immediate transplant<br>Group 4: static warm storage (6 hrs) + transplant<br>Group 5: flush w/ HTK (10 mL) + SCS (4 °C, 6 hrs) + transplant | 5-10 | 65 | 100% of groups 2 and 3, 80% of group 5, and 0% of group 4 limbs survived to 12 weeks<br>Group 4: higher muscle injury score than group 5<br>Group 5: lowest CMAP amplitude and maximal twitch force of all groups, extensive muscle degeneration and loss of organization with foci of regeneration. |

Note: DMSO = dimethyl sulfoxide; RL = Ringer's lactate solution; UW = University of Wisconsin solution; SCS = static cold storage; Gold standard = flushing with UW and subsequent SCS with UW; HTK = histidine-tryptophan-ketoglutarate solution; CMAP = compound muscle action potential. Details regarding time, temperature, amount, or identities of solutions not listed were not reported by the study.

Rinker et al cryopreserved grafts at -140°C for two weeks and showed a viability index of 91.9% after two weeks. However, upon transplantation, all but one graft in the cryopreserved group failed by venous thrombosis, compared to 100% survival in freshly transplanted grafts. The authors attribute the failure by venous thrombosis to IRI.^22^

Liu et al preserved hemifacial flaps with UW solution beyond the gold standard interval for 24, 36, and 48 hours to establish an upper time limit for VCA viability.^25^ The study demonstrated a decrease in CMAP amplitude and extended latency from baseline across all groups. The corneal reflex in these hemifacial flaps recovered after 6 months for groups undergoing 24 hours of SCS and after 9 months for groups undergoing 48 hours of SCS. In separate tissue analyses, the viability of muscles and nerves was found to have decreased most quickly while skin remained viable for longer. Based on the lack of statistically significant differences in overall graft viability between 12 and 24 hours, the authors chose 24 hours as the time limit for the gold standard for preservation.^25^

Amin et al evaluated the addition of a Ringer’s lactate solution flush after the standard SCS protocol to remove leukocytes, however there was no significant difference in leukocyte levels in the venous effluent, aside from a decrease in CXCL-8 compared to a no flush control.^12^

Gok et al 2020 studied the effect of a post-harvest flush on muscle viability comparing graft changes across various groups: naïve, sciatic nerve transection and repair, immediate transplant, and static warm storage with transplant.^23^ The investigators found that 80% of SCS hindlimbs survived to the 12-week endpoint but with a significant muscle injury score of 73% and decreased CMAP amplitude and contractile force compared to baseline. Flaps preserved under static warm storage did not survive and had a high muscle injury score.^23^ A summary of the four study protocols is shown in **Table 3**.

## Discussion

In this scoping review, we identified and analyzed 13 studies on pre-treatments in VCA, with a focus on graft viability. We grouped PTs into three categories: solution-based alterations to the gold standard, *ex vivo* perfusion, and other techniques. PTs altering UW solution focused on attenuating destructive changes post-transplantation, while the other novel methods group primarily targeted ischemia-specific destruction. *Ex vivo* perfusion studies attempted to minimize all harm to the graft to demonstrate feasibility of this technique in VCA. Below, we elaborate on themes and considerations across these developing PT methodologies, highlight their potential uses, and suggest future studies needed for clinical translation.

### Solution-based Alterations to the Gold Standard

As is the overarching goal of preservation, the choice of pre-treatment solution aims to maintain stability of the graft environment during ischemia time to attenuate any future tissue damage. Solution choice should consider buffering pH changes, providing oxygen, maintaining energy stores and osmotic balance, preventing clotting and buildup of toxic metabolites, among many other considerations. Solutions should not harm the graft, develop a precipitate, or readily extravasate into tissues.

Replacing UW solution in the gold standard protocol with UW solution with H_2_S or Perfadex yielded encouraging results. Of the reviewed methods, these PTs would be the easiest to implement, as they require no additional technical considerations. In a 2014 VCA study, H_2_S-treated grafts had statistically significantly lower levels of creatine kinase, lactate dehydrogenase, aspartate transaminase, and TNF-α after three hours of ischemia compared to UW solution alone.^30^ Fries at al also demonstrated a delay in the onset of acute rejection with the addition of H_2_S emphasizing the potential to reduce early post-transplantation inflammation. Targeted delivery of H_2_S would be a unique, non-toxic route for enhancing the efficacy of immunosuppression. However, this study evaluated the shortest preservation time at only three hours.^18^ Further studies are necessary to determine the efficacy of H_2_S over longer PT intervals.

Perfadex is a low-potassium, dextran-based solution that has been used in lung transplantation and is being explored an as option in preservation for VCA.^31–33^ This solution showed promise in reducing IRI markers in the Rostami et al study, which also highlighted tissue-specific vulnerability to IRI.^20^ The increased susceptibility of muscle to ischemia is validated by the literature and other papers in this review, which describe edema, inflammation, myofibril disorganization, non-homogeneous necrosis, among other maladaptive alterations in tissue architecture.^23,24,27–29,34^ IRI may compromise graft viability by disrupting laminin and CD31 indicating reduced vascular integrity jeopardizing the primary mode of communication between the graft and recipient.^20,9^ In contrast, in all reviewed studies, skin and bone were shown to be resilient against ischemic damage.^20^ Given the heterogeneous tissue setting, the success of VCA relies on careful evaluations of each tissue type, as shown by Liu et al.^25^

### *Ex vivo* Perfusion

Machine perfusion has gained considerable interest in the VCA community in recent years due to its ability to continuously deliver nutrients, buffer cells, reduce inflammatory modulators, and remove toxins.^11^ Historic success in SOT validates perfusion as a topic of investigation for VCA research.^17,35^ Various renal transplantation studies have reported reduced vascular resistance after reperfusion, decreased risk and duration of delayed graft function, and higher graft survival rates.^36–38^ Based on the evidence, the U.S. Food and Drug Administration approved the first *ex vivo* liver perfusion device in 2021.^39^ These advances suggest potential for similarly improved preservation potentially resulting in positive patient outcomes in VCA.

The studies in our review yielded promising results of perfusion for muscle preservation in particular. As muscle is the dominant tissue component of VCAs, its preservation is paramount in our ability to enhance functionality by optimizing motor recovery. Unfortunately, given its high susceptibility to ischemia, muscle viability begins to deteriorate after only four hours post-harvest.^7^ The decrease in muscle function was shown in both the SCS control groups in the perfusion studies and in the study by Liu et al with 24 hours of SCS. After 24 hours of perfusion, Ozer et al 2016 demonstrated preserved muscle contractility similar to a non-operated control 12 hours after transplant.^21^ Ozer et al 2015 and Constantinescu et al showed similar success.^20,27^ For clinical translation, future studies should extend the observation timeline to further characterize muscle viability through IRI and acute rejection intervals.

Flow parameters and perfusate choice may play a role in fluid buildup and should be thoughtfully considered prior to use. Graft weight gain, attributed to fluid buildup, was seen in all seven perfusion studies with weights up to 150% of baseline.^26^ Burlage et al observed decreased weight gain with the addition of HBOC-201 and PEG to skeletal muscle media, substances that help retain osmotic pressure within the vasculature and reduce endothelial leakage.^24^ Another concern with closed circuit perfusion are the complexity and cost of equipment.^40^ Toxic metabolite buildup requires continuous monitoring and management by an active researcher, adding to the resource demand with this PT method. These technical considerations may render perfusion inappropriate for some institutions necessitating the development of strategies to mitigate barriers to wider implementation.

### Other Techniques

These final studies presented innovative strategies for preserving VCAs, including cryopreservation for long-term storage and adding a post-SCS flush intended to reduce the inflammatory burden of the graft. Other studies aimed to establish a preservation time limit with the gold standard and its impact on muscle viability and function.

These studies emphasize an important theme in VCA transplantation: a discordance in defining viability by histological assessment (pre- or post-transplantation) vs. by clinical outcomes. After two weeks of cryopreservation, grafts showed high histological viability, yet 90% of grafts failed after transplantation.^22^ Concurrently, in the Gok et al studies, transplanted grafts preserved with SCS showed significant muscle injury, yet most grafts survived with preserved muscle function.^23,29^ In the Fries et al study previously discussed, there was no histological distinction between H_2_S- and UW-treated grafts, yet the H_2_S-treated grafts demonstrated delayed rejection.^18^ A 2019 study found that clinical response in acute graft-versus-host-disease correlated with prognosis while histology did not.^41^ Together, these findings underscore the importance of the post-transplantation clinical evaluation of VCAs in PT experimentation. This suggests prioritization of clinical observations over histological findings in cases of discrepancy and the need for tailoring histological evaluation of rejection, such as the Banff criteria, to VCA specifically.^42^

Regardless, these results show promise, particularly the high graft viability despite extended cryopreservation storage times. With the increasing utilization of cryopreservation for *in vitro* fertilization, including gamete and embryo preservation, cryopreservation remains a relevant topic of investigation.^43,44^ Use of heparin in this protocol (**Table 3**) is a potential future direction to explore in cryopreservation research as grafts failed by thrombosis.

To determine a preservation time limit for the gold standard protocol, Liu et al used the longest follow-up period at nine months, when the last canine corneal reflex recovered.^25^ As VCAs contain primarily muscle tissue, follow-up times must be sufficiently long to adequately characterize the recovery of this vital portion of the graft and analyze the degree of functional recovery using different PT methods. At three and six months, the Gok et al 2020 and Liu et al studies, evaluating SCS preservation, showed diminished muscle response on electromyography.^23,25^ Finally, although immunological impact is out of the scope of this review, Amin et al made a unique inquiry into the impact of leukocyte burden on graft injury.^12^ Given the proposed link between inflammation during ischemia and IRI and rejection, future research in this area is warranted.

### Limitations

This study has limitations common to retrospective and review studies; notably, the findings analyzed here rely on the fidelity of data reported in the included studies. Studies may also have been erroneously excluded in the initial screening process. While we could not conduct statistical heterogeneity testing, methods and reporting were considerably varied, and we could not perform re-analysis of PT data reported across multiple studies. Given the demanding nature and complexity of VCA, the primary limitation for this review is that clinical studies remain scarce, restricting the breadth and depth achievable by systematic reviews at this time. The limited number of included studies, as well as the small sample sizes within these studies, also diminish the extent to which definitive conclusions can be drawn. Despite these limitations, this review provides a glance into the current state of PT in VCA and identifies areas for further exploration.

## Conclusion

This study of pre-treatment modalities found a variety of promising techniques in the vascularized composite tissue preservation setting. As VCA is relatively novel in the field of transplantation, many of the preservation techniques have been informed by the SOT literature. However, owing to its tissue heterogeneity, VCA presents a unique list of challenges, stifling direct translation of SOT and preclinical to clinical work. Overall, the PTs analyzed showed success with limited clinical translation. With increasing time and resources being invested into research on VCA preservation, the donor and recipient pools should expand concomitantly. This will further validate the safety and efficacy of these solutions and drive the establishment of federal regulatory protocols and a standard of care in VCA. The results of this scopimh review demonstrate the promise of future research, particularly machine perfusion.

---

**Supplemental Table 1.**
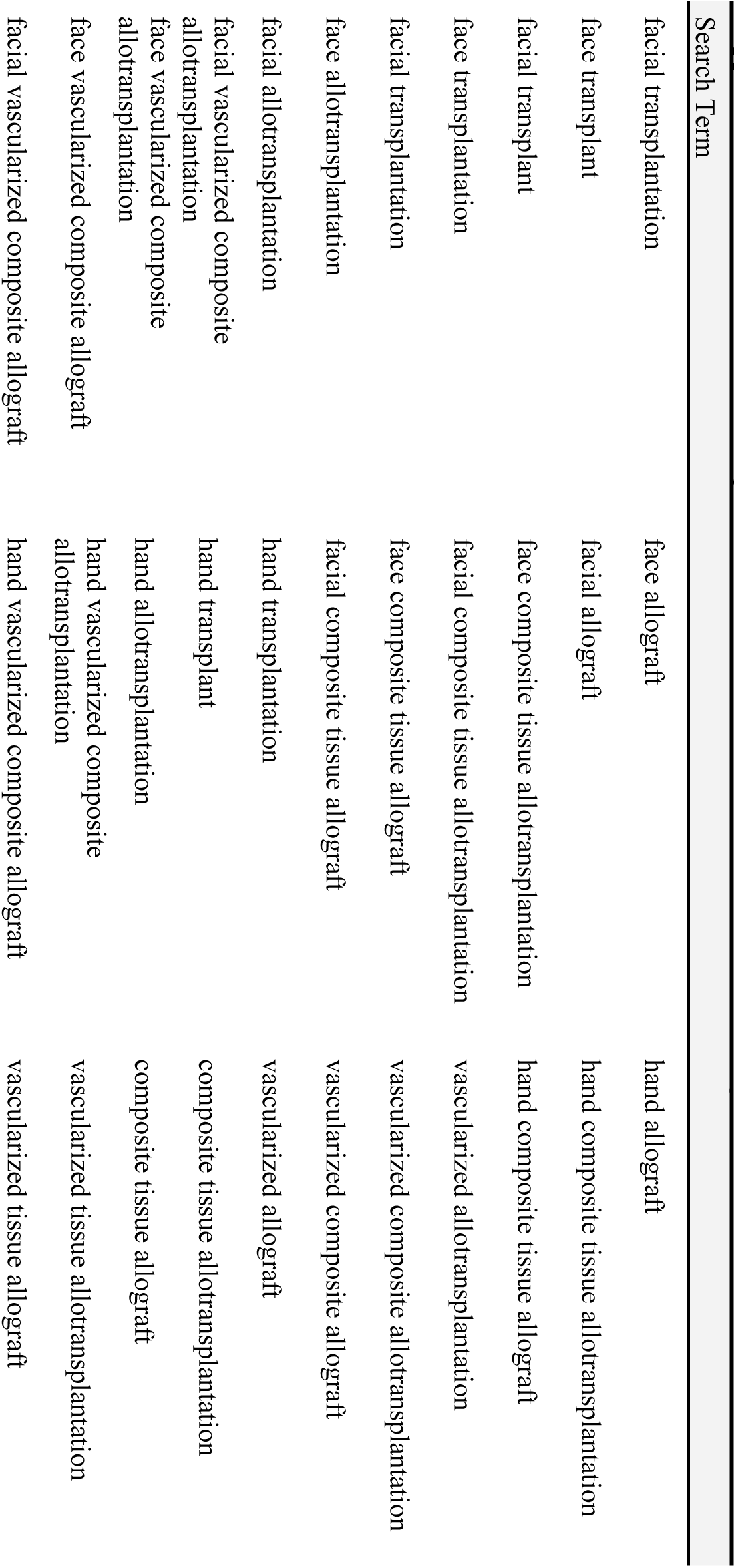
Article screening search terms.

**Supplemental Table 2.**
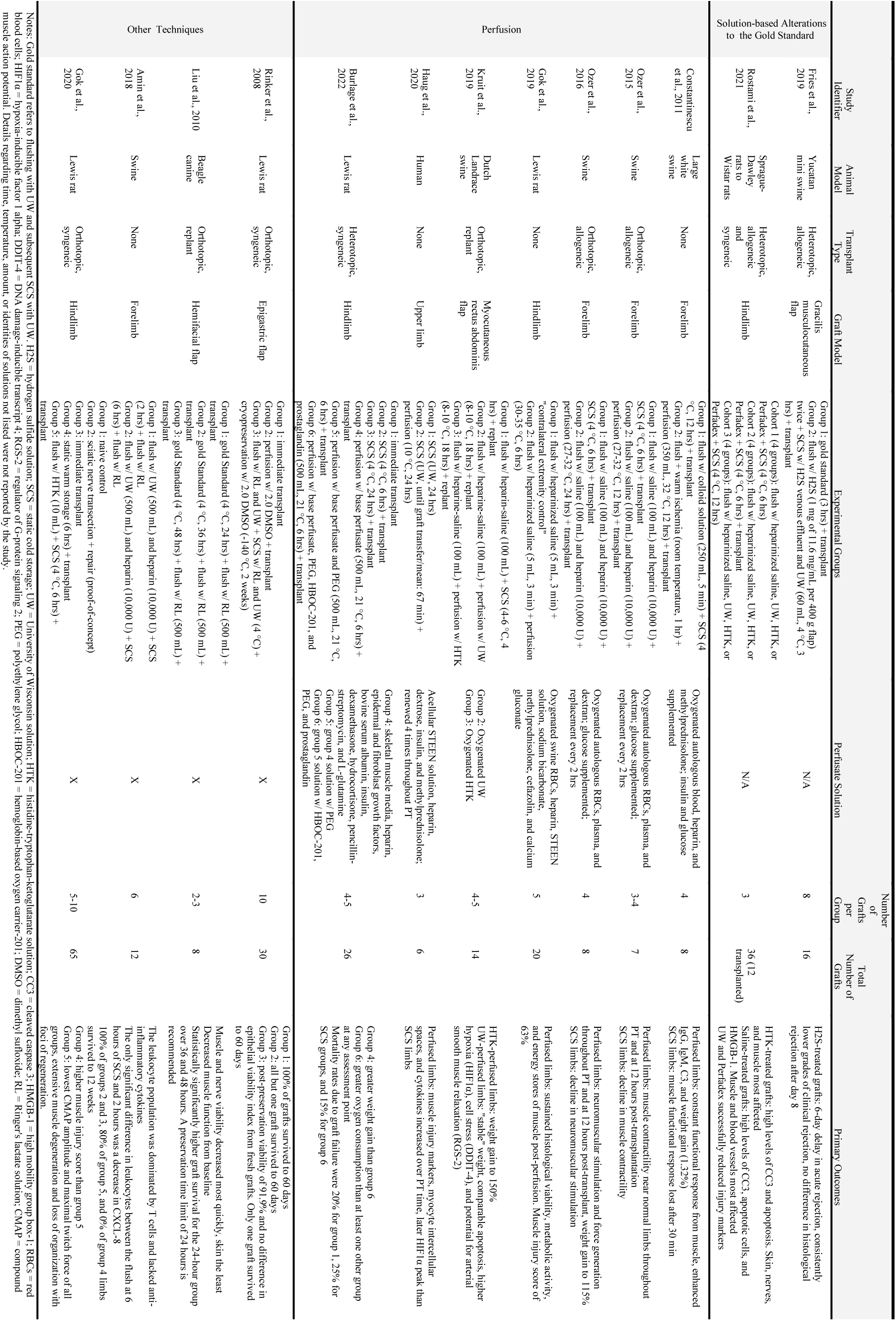
Characterizations of all pre-treatment experimental protocols reviewed.

